# A Novel Family of Genomics Islands Across Multiple Species of *Streptococcus*

**DOI:** 10.1101/065920

**Authors:** Jing Wang, Changjun Wang, Weizhong Feng, Youjun Feng, Liming Zhi, Wenjuan Li, Yi Yao, Shibo Jiang, Jiaqi Tang

## Abstract

The genus *Streptococcus* is one of the most genomically diverse and important human and agricultural pathogens. The acquisition of genomic islands (GIs) plays a central role in adaptation to new hosts in the genus pathogens. The research presented here employs a comparative genomics approach to define a novel family of GIs in the genus *Streptococcus* which also appears across strains of the same species. Specifically, we identified 9 *Streptococcus* genomes out of 67 sequenced genomes analyzed, and we termed these as 15bp *Streptococcus* genomic islands, or 15SGIs, including i) insertion adjacent to the 3’ end of ribosome *l7/l12* gene, ii) large inserts of horizontally acquired DNA, and iii) the presence of mobility genes (integrase) and replication initiators. We have identified a novel family of 15SGIs and seems to be important in species differentiation and adaptation to new hosts. It plays an important role during strain evolution in the genus *Streptococcus*.

## Introduction

Several studies have suggested that recombination might be the key factor in the adaptation of pathogens to new niches, antibiotics or immune systems (Sentchilo *et al.*2009; Marri *et al.* 2010). In most bacterial species, horizontal gene transfer (HGT) and the acquisition of foreign DNA are widely recognized as the mechanisms by which bacteria adapt to changing environments (Lefébure *et al.* 2007). The horizontally acquired DNA fragments, such as plasmids, bacteriophages, transposons, integrative and conjugative elements, and genomic islands (GIs), allow bacteria to instantly obtain a range of genetic traits that may increase fitness under different environmental conditions (Juhas *et al.* 2009), and even under extreme environmental conditions, to defeat host immune system defenses (Tuanyok *et al.* 2008). GIs are classified on the basis of the different functions they perform, particularly the encoding of major gene proteins, such as those involved in metabolism, antimicrobial resistance, symbiosis and pathogenicity (i.e., pathogenicity islands, PAI) (Dobrindt *et al.* 2004). Specifically, if we can detect the presence of multiple sequenced closely related *Streptococcus* genomes that inhabit different niches, we will not only be able to understand the pattern of their genomic evolution but we will gain an excellent dataset giving us insight into the role of species-specific genomic islands. Nine genomes of streptococcus belong to four species which have previously been analyzed: *Streptococcus pneumoniae, Streptococcus agalactiae, Streptococcus suis* and *Streptococcus uberis.*

*Streptococcus pneumoniae* is the leading cause of community acquired pneumonia, sepsis, and meningitis that annually results in over a million deaths worldwide. Analysis of publicly available genomes of *Streptococcus pneumoniae* has presented a few GIs associated with virulence and antimicrobial resistance, for example, pneumococcus pathogenicity island 1(PPI1) and the resistance island encoding eight resistance genes (Brown *et al.* 2004).

*Streptococcus agalactiae* is a Lancefield’s group B Streptococcus (GBS). It is the leading cause of meningitis and sepsis in newborns. Additionally, this pathogen is the cause of serious infections in immunocompromised adults. Clinical manifestations of infection include urinary tract and soft tissue infections, as well as life-threatening sepsis and meningitis (Glaser *et al.* 2002). It is also implicated as a pathogen in cases of mastitis in cows (Petzer *et al.* 2009). Analysis of the genomes of two strains, NEM316 and 2603VR, resulted in the annotation of 14 putative PAI-like structures, but without conclusive data to confirm their association (Herbert *et al.* 2005). Two novel GI families with pilus-like coding and containing a two-component signal transduction system were reported separately in 2006. Based on the islands which code for pilus-like structures, important virulence factors and, hence, potential vaccine candidates were proposed (Rosini *et al.* 2006). Still another PAI-like structure contained genes homologous to the sensor histidine kinase gene and the DNA-binding response-regulator gene of *Streptococcus pneumoniae* and the bac gene in *Streptococcus agalactiae* (Dmitriev et al. 2006).

*Streptococcus suis* is responsible for a variety of diseases in pigs, including meningitis, septicemia, arthritis, and pneumonia. It is also a zoonotic pathogen that causes occasional cases of meningitis and sepsis in humans, but it has also recently been implicated in outbreaks of streptococcus toxic shock syndrome by our group. In 2007, we descried an 89-kb PAI-like structure in the highly virulent Chinese strains 05ZYH33 and 98HAH12. This 89-kb PAI-like structure includes a specific two-component signal transduction system which can regulate the virulence of strain 05ZYH33 (Chen *et al.* 2007; Li *et al.* 2008).

*Streptococcus uberis,* a Gram-positive bacteria pathogen responsible for a significant proportion of bovine mastitis in commercial dairy herds, colonizes in multiple body sites of the cow, including the gut, genital tract and mammary gland. Comparative genomics analysis revealed *Streptococcus uberis* to be more similar to *Streptococcus agalactiae* than either *Streptococcus pneumoniae* or *Streptococcus suis.* In comparison with other phylogenic streptococci, a lower number of mobile genetic elements were displayed, but there were bacteriophage-derived islands, and a putative GI was identified (Ward *et al.* 2009).

In this study, we identified a novel 15 SGI family that is distributed among the genus of streptococcus belonging to the above four species, and we analyzed the new structure of this GI family. The presence of multiple gene content closely related to the GIs of these genomes suggests that horizontal gene transfer (HGT) plays an important role in the evolution of these *Streptococcus* species and subspecies adapting to new niches.

## Methods

### Streptococcus strains, media, and culture condition

Analysis of distribution of 15SGI was performed on clinical streptococcus isolates, including 10 *Streptococcus pneumoniae* strains, 10 *Streptococcus agalactiae* strains, one *Streptococcus mitis,* one *Streptococcus sanguis* and 17 *Streptococcus suis* strains, which consist of 3 strains from sporadic cases of meningitis in China, 12 strains from Europe, and the strains 05ZYH33 and 03HAS68. All strains were cultured at 37°C on Columbia agar supplemented with 5% sheep blood or in Todd-Hewitt broth supplemented with 0.5% yeast extract (THY).

### PCR analysis of GIs

Primers for amplifying these specific sequences were designed with Primer Premier 6 (Table 4). PCR experiments used for detecting 15bp repeat sequence from l7/l12 gene terminal and adjacent integrases were preformed in a final volume of 50μl. After an initial denaturation step (3min at 94°C), a 32-cycle PCR was performed (denaturation 30s at 94°C, annealing 30s at 52°C, extension at 72°C for 1min), followed by a final extension step (5min at 72°C).

**Table 4.**
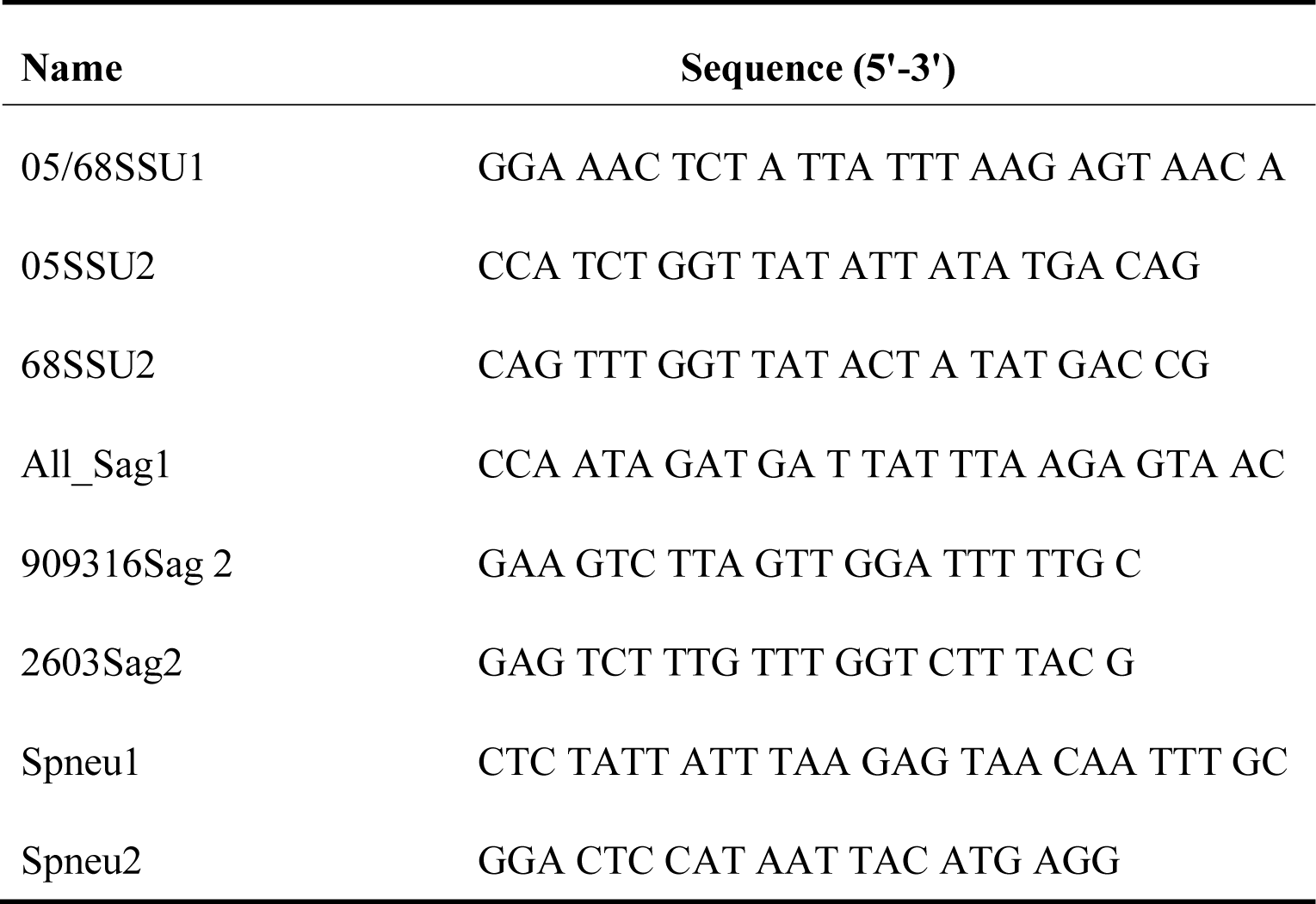
Primers used in this study.

### Genomic dataset

All the sequences of this study involved 54 streptococcus complete genomes, and the annotations were downloaded from the National Center for Biotechnology Information (NCBI) database (http://www.ncbi.nlm.nih.gov/genomes/genlist.cgi?taxid=2&type=0&name=Complete%20Bacteria). Throughout the analysis of 15bp repeat sequences and mobile elements, we focused on the analysis of 9 reference bacterial strains from four different species of streptococcus, including *Streptococcus pneumoniae, Streptococcus agalactiae, Streptococcus suis* and *Streptococcus uberis* (Table 1). Analyses of G+C content of genomes were shown by CLC Genomics Workbench 3 software.

**Table 1.** Information about the nine streptococcus strains and 15bp genomic islands.

| **Species** | **Genomes** |  |  | **GIs** |  |  |
| --- | --- | --- | --- | --- | --- | --- |
|  | **Strain** | **Source** | **Rsfseq**<br>**accession**<br>**No.** | **Coordinates** | **GI**<br>**Size** | **GC%**<br>**deviation** |
| *S.uberis* | 0140J | GenBank | NC_012004 | 768595..783713 | 15119 | 0.32 |
| *S.pneumoniae* | ATCC700669 | GenBank | NC_011900 | 1259646..1327358 | 67713 | -5.23 |
| *S.pneumoniae* | CGSP14 | Genbank | NC_010582 | 1207719..1288815 | 81097 | -5.5 |
| *S.suis* | 05HAS68 | GenBank | CP_002007 | 1148502..1261166 | 112665 | -281 |
| *S.suis* | 05ZYH33 | GenBank | NC_009442 | 872952..961833 | 88882 | -4.28 |
| *S.suis* | BM407 | GenBank | NC_012926 | 1005780..1086173 | 80394 | -3.95 |
| *S.agalactiae* | 2603V/R | GenBank | NC_004116 | 1256620..1311411 | 54792 | 2.68 |
| *S.agalactiae* | NEM316 | GenBank | NC_004368 | 1361778..1423551 | 61774 | 2.32 |
| *S.agalactiae* | A909 | GenBank | NC_007432 | 1309001..1319073 | 10073 | -1.08 |

### Comparative genomics and prediction of GI

Each genomic sequence comparison was preformed from the NCBI BLAST program, which is available in the BioEdit Sequence Alignment Editor. The results were visualized using ACT (Artemis Comparison Tool) release 8 based on Java(TM) web start launcher. The files of all 15SGIs were obtained from GenBank by Vector NTI 10 and visualized by using CLC Genomics Workbench 3 software. The figures were configured using ACT and CLC Genomics Workbench exported graphics (Fig. 2).

**Figure 1.**
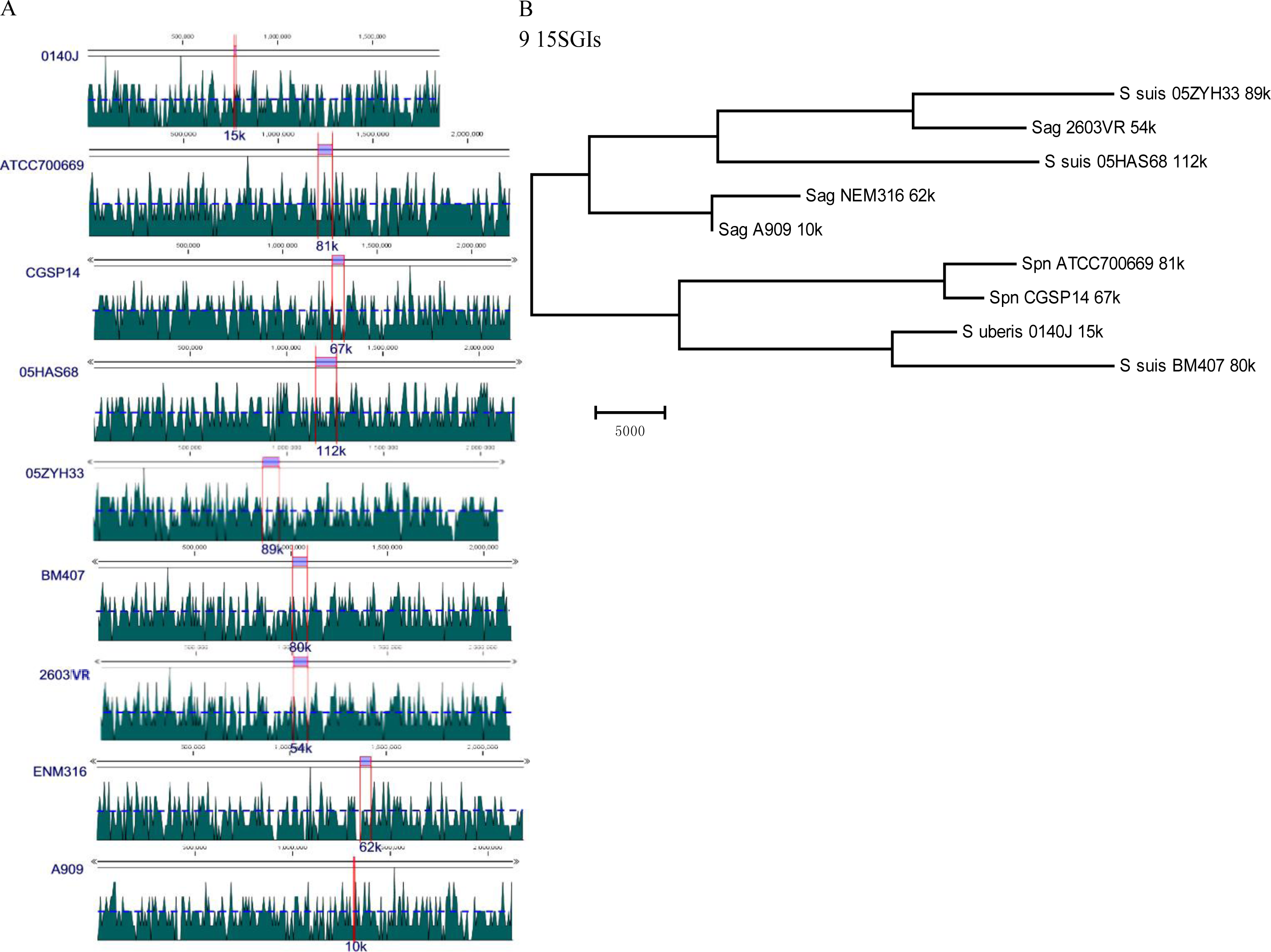
Analysis of GC content and phylogenetic comparison of 15SGIs. A. GC contents of 9 *Streptococcus* genomes. The area chart representing the GC percentage of the genomes. The red box represents the 15SGI segments in strains, respectively. B. Phylogenetic comparison of 15SGIs. The tree was constructed with 15SGI sequences of different *Streptococci* by using MEGA 4. Images represent genetic distance as number of nucleotide substitutions.

**Figure 2.**
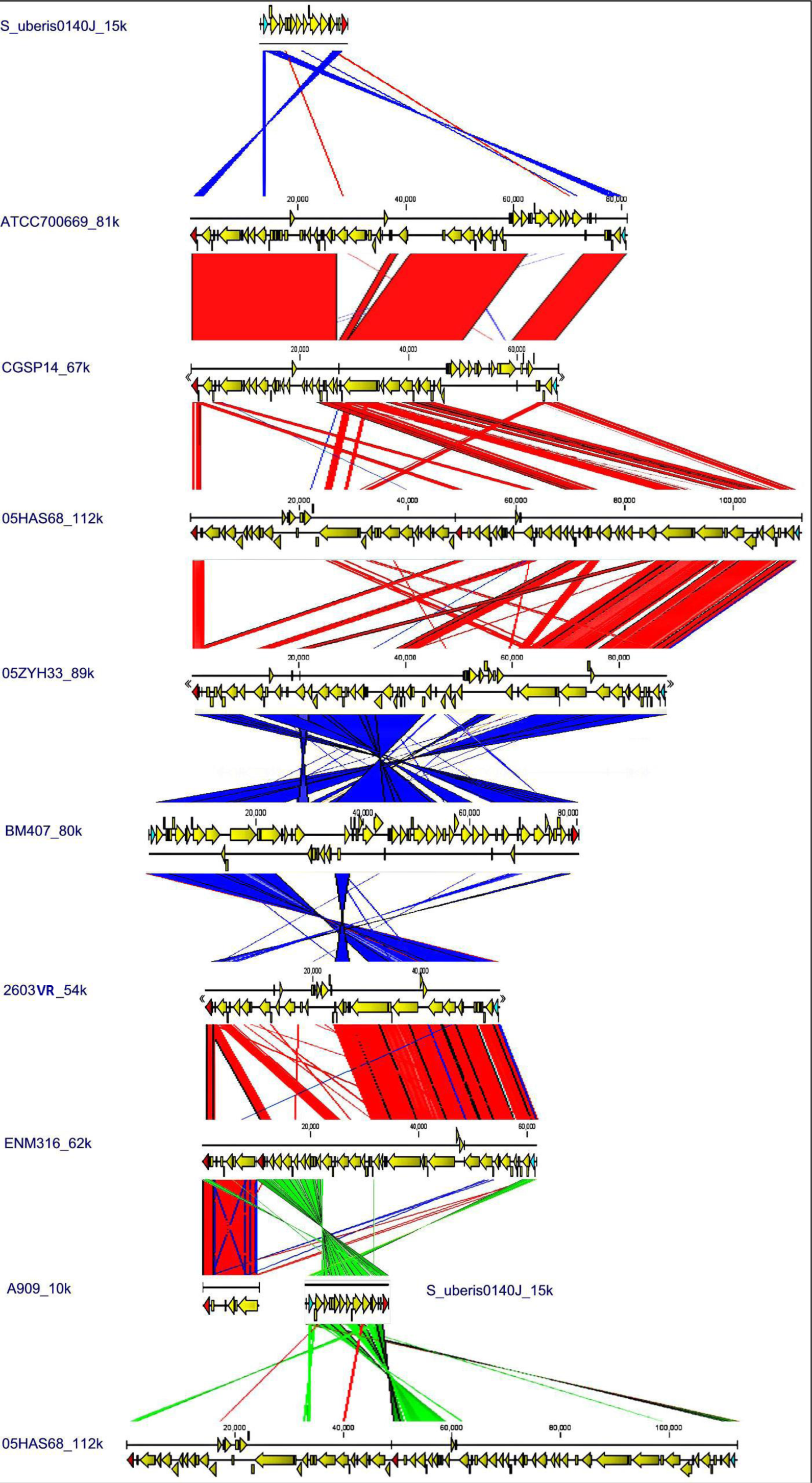
Comparisons of the 15SGI regions from 9 *Streptococcus* strains. Pairs of the 15SGI region between two strains were generated by TBLASTN matches and linear comparison using the Artemis Comparison Tool (ACT). The ORFs of 15SGIs were displayed using CLC Genomic Workbench. Red or green lines indicate TBLASTN matches between genomes in the same direction; blue lines indicate TBLASTN matches between genomes in opposite directions.

### Identification of genes

The protein sequences from all 15SGIs were compared using BlastP with matrix BLOSUM62 with an expectation value set at 1.0E-20 and an additional criterion of match length set at 75% of the query sequence.

### Phylogenetic analysis

The evolutionary history of 9 15SGIs was inferred using the Maximum Parsimony method. The most parsimonious tree with length = 162406 is shown. The consistency index is 0.848879 (0.781436), the retention index is 0.646868 (0.646868), and the composite index is 0.549113 (0.505486) for all sites and parsimony-informative sites (in parentheses). The MP tree was obtained using the Close-Neighbor-Interchange algorithm with search level 3 in which the initial trees were obtained with the random addition of sequences (10 replicates). Branch lengths were calculated using the average pathway method and are in the units of the number of changes over the whole sequence. They are shown next to the branches. The tree is drawn to scale. All alignment gaps were treated as missing data. There were a total of 115353 positions in the final dataset, out of which 49497 were parsimony informative.

The evolutionary history of 11 intregrases and 8 replicator initiators of 15SGIs were inferred using the UPGMA method. The bootstrap consensus tree inferred from 500 replicates is taken to represent the evolutionary history of the taxa analyzed. Branches corresponding to partitions reproduced in less than 50% of bootstrap replicates are collapsed. The percentage of replicate trees in which the associated taxa clustered together in the bootstrap test (500 replicates) is shown next to the branches. The tree is drawn to scale, with branch lengths in the same units as those of the evolutionary distances used to infer the phylogenetic tree. The evolutionary distances were computed using the Poisson correction method and are in the units of the number of amino acid substitutions per site. All positions containing gaps and missing data were eliminated from the dataset (complete deletion option). There were a total of 335 and 255 positions in the final dataset. All phylogenetic analyses were conducted in MEGA4 (Tamura *et al.* 2007).

### Homology modeling of the integrase core domain

Amino acids sequences of 11 integrases from 9 15SGIs were submitted to a fully automated protein structure homology-modeling server of SWISS-MODEL (Arnold *et al.* 2006 http://swissmodel.expasy.org/) and a server for prediction of protein-DNA interaction sites of DNABindR (http://turing.cs.iastate.edu/PredDNA/predict.html), respectively. Structural representations of the integrase models were created with the free molecular-graphics program PyMOL.

## Results

### Distribution and structural features

In all, we analyzed 67 genomes of *Streptococcus* across 12 different genera found on public databases, including those of the NCBI and the Sanger Institute. We then made a side-by-side comparison using the Artemis Comparison Tool (ACT), which enabled genomic alignment and visualization of BLASTN results. Intriguingly, 9 reference strains of three different genera displayed structurally similar GIs (Table 1). Based on the fact that insertion of GIs into the bacterial chromosome is often a site-specific event, these GIs further shared two 15bp direct repeats in two respective attachment sites, attL and attR, of *Streptococcus* genomes, which we termed the 15bp *Streptococcus* genomic island (15SGI).

Structurally, 15SGIs share the 15bp direct repeats in common, but they share variant G+C contents with their host bacteria. Large arrays of proven and putative GIs or PAIs from Gram-positive bacteria generally showed a low G+C content, while a few showed higher G+C content. In the 9 reference streptococcus strains, G+C content of 15SGI in *Streptococcus pneumoniae* ATCC700669 and CGSP14, as well as *Streptococcus suis* 05ZYH33, 05JYS68 and BM407, is significantly lower than the overall G+C content of the host. While G+C content of 15SGI in *Streptococcus agalactiae* 2603V/R and NEM316 is higher, it is the same as both *Streptococcus agalactiae* A909 and *Streptococcus uberis* 0140J (Table 1, Fig. 1). Our results also indicated that the GIs are mosaic and that some genes may have been acquired recently. New gene acquisitions may have atypical sequence characteristics that are maintained over a relatively short time frame. For example, in *Streptococcus uberis* 0140J, the 15SGI locus was changed, indicating that an unknown putative GI was inserted between 15SGI and *17/112.* The putative GI, which was predicted by Ward et al. (Ward *et al.* 2009), was about 10kb with a G+C content of 34.79%, compared to 15SGI with a mean of 36.95% and the overall *Streptococcus uberis* 0140J genome mean of 36.63%. The reading frame position-specific GC usage indicated that the DNA fragment may have been more recently acquired than the 15SGI. The deviation of G+C content is not obvious from the 15SGIs of *Streptococcus agalactiae* strain A909, although it is, perhaps, smaller in relation to the others.

Also, in these 15SGIs, we find the presence of mobility genes which 1) encode a site-specific recombinase/integrase to the right of attL and 2) encode the replication initiator located to the left of attR. Distribution analyses of 15SGI with bacterial genomes downloaded from GenBank indicated that they were not related to the serotype of *Streptococcus* intra-subspecies. In contrast, distribution analyses of 15SGI with clinical isolates of *Streptococcus* by PCR showed that it has more widespread distribution in different *Streptococcus* species, such as *Streptococcus mitis.*However, 15SGI integration did not occur in the strains of *Streptococcus suis* from 3 strains of sporadic cases with meningitis in China, 12 strains from Europe, and 10 isolates of *Streptococcus pneumoniae.*

### Tandem accretion and deletion by site-specific recombination

To analyze the structural features of nine 15SGIs from *Streptococcus,* two 15SGI tandem arrays emerged from integration at the end of the *17/U2* gene in the chromosome of *Streptococcus agalactiae* strain 2603V/R and *Streptococcus suis* strain 05ZYH33, respectively. The structure of the first 15SGI shows complete integrity, as it was directly linked to the 3’ end of *17/U2* gene based on the 15bp integration hotspot; however, the *replicative initiator A* gene was deleted from the second 15SGI. This phenomenon may reflect a complex evolutionary history of the GI, possibly enabling it to be mobile, as represented by the 48-kb, 64-kb or 112-kb DNA segments of *Streptococcus suis* strain 05HAS68 or the 10-kb, 52-kb or 62-kb DNA segments of the *Streptococcus agalactiae* strain NEM316. We know that GI insertion is time dependent. Therefore, by comparing the deviation of G+C content between GI and host chromosome could give a good indication of the time of insertion. Specifically, since the GC content of 48-kb and 64-kb is 36.45% and 39.83% from *Streptococcus suis* strain 05HAS68, respectively, in comparison with its overall genome mean of 41.18%, it appears that the 48-kb fragment was integrated later than the 64-kb fragment. Similarly, the DNA fragment of 10-kb and 52-kb display 34.58% and 38.61% GC content from *Streptococcus agalactiae* strain NEM316, respectively, in comparison with its overall genome mean of 35.63%; therefore, it appears that the 52-kb fragment was integrated later than the 10-kb fragment. Interestingly, compared with the 10-kb 15SGI from *Streptococcus agalactiae* strain A909 and the DNA fragment of 10-kb of 62-kb 15 SGI of *Streptococcus agalactiae* strain NEM316 is highest identify in the ACT alignment and share no containing *replicative initiator A* gene at their 3’ region (Fig. 2). In addition, from recent unpublished studies, we have found that the 89-kb 15SGI could be excised from the chromosome of *Streptococcus suis* strain 05ZYH33 and form circular intermediates. We therefore infer two 15SGIs between *Streptococcus agalactiae* strain NEM316 and A909 as having the same the foreign genetic source material, but the second 15SGI (52-kb fragment) excised from the chromosome of *Streptococcus agalactiae* strain A909.

### Gene contents and potential functional roles

As a rule, GIs often carry genes that reflect a selective advantage to the host bacterium in a specific environment. Among the nine 15SGIs, this study revealed the presence of genes belonging to the genus *Streptococcus* across 4 species, carrying only a few uniquely specific genes for the 15SGI of strain and species, but the large across species genes content were emerged by comparing with protein sequences of genes coding from all the 15SGIs each other (Fig. 3). Our previously studies indicated that GI-typeT4SS genes are required for 89k PAI-like of *Streptococcus suis* 05ZYH33 strain (Zhao *et al.* 2011). 7 of 9 15SGIs carry 4 genes of GI-type T4SS and identity of them are more than 60% (Fig 2., Table 2). These data indicate that 15SGIs provide *Streptococcus* with a readily available novel gene pool that supplies additional physiological properties from not only one identified biological source, but also multiple sources which help it to exploit new niches.

**Table 2.**
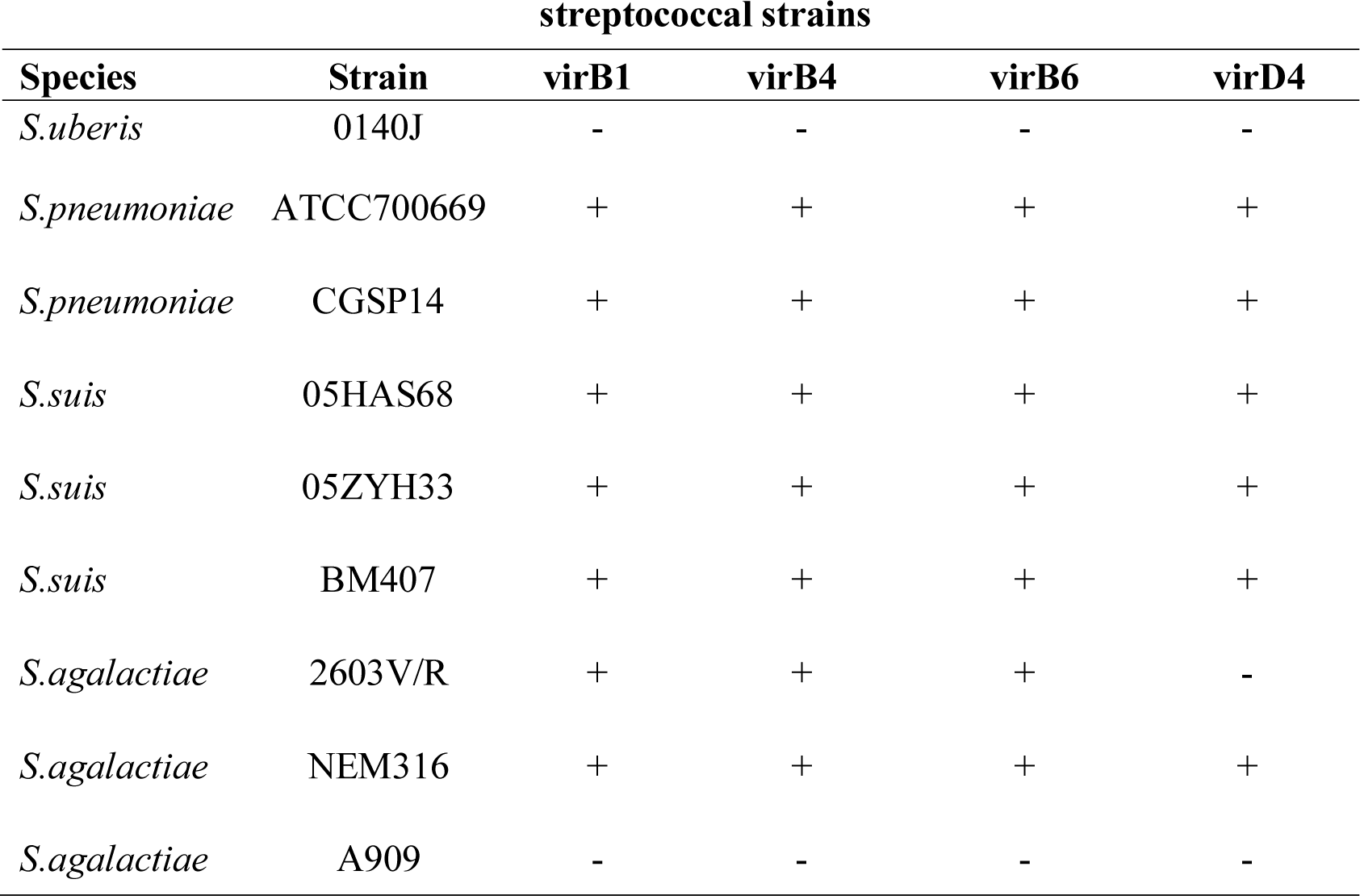
The distribution of genes of GI-type IV secretion systems in 15SGI from.

| **Species** | **Strain** | **virB1** | **virB4** | **virB6** | **virD4** |
| --- | --- | --- | --- | --- | --- |
| *S.uberis* | 0140J | - | - | - | - |
| *S.pneumoniae* | ATCC700669 | + | + | + | + |
| *S.pneumoniae* | CGSP14 | + | + | + | + |
| *S.suis* | 05HAS68 | + | + | + | + |
| *S.suis* | 05ZYH33 | + | + | + | + |
| *S.suis* | BM407 | + | + | + | + |
| *S.agalactiae* | 2603V/R | + | + | + | - |
| *S.agalactiae* | NEM316 | + | + | + | + |
| *S.agalactiae* | A909 | - | - | - | - |

**Figure 3.**
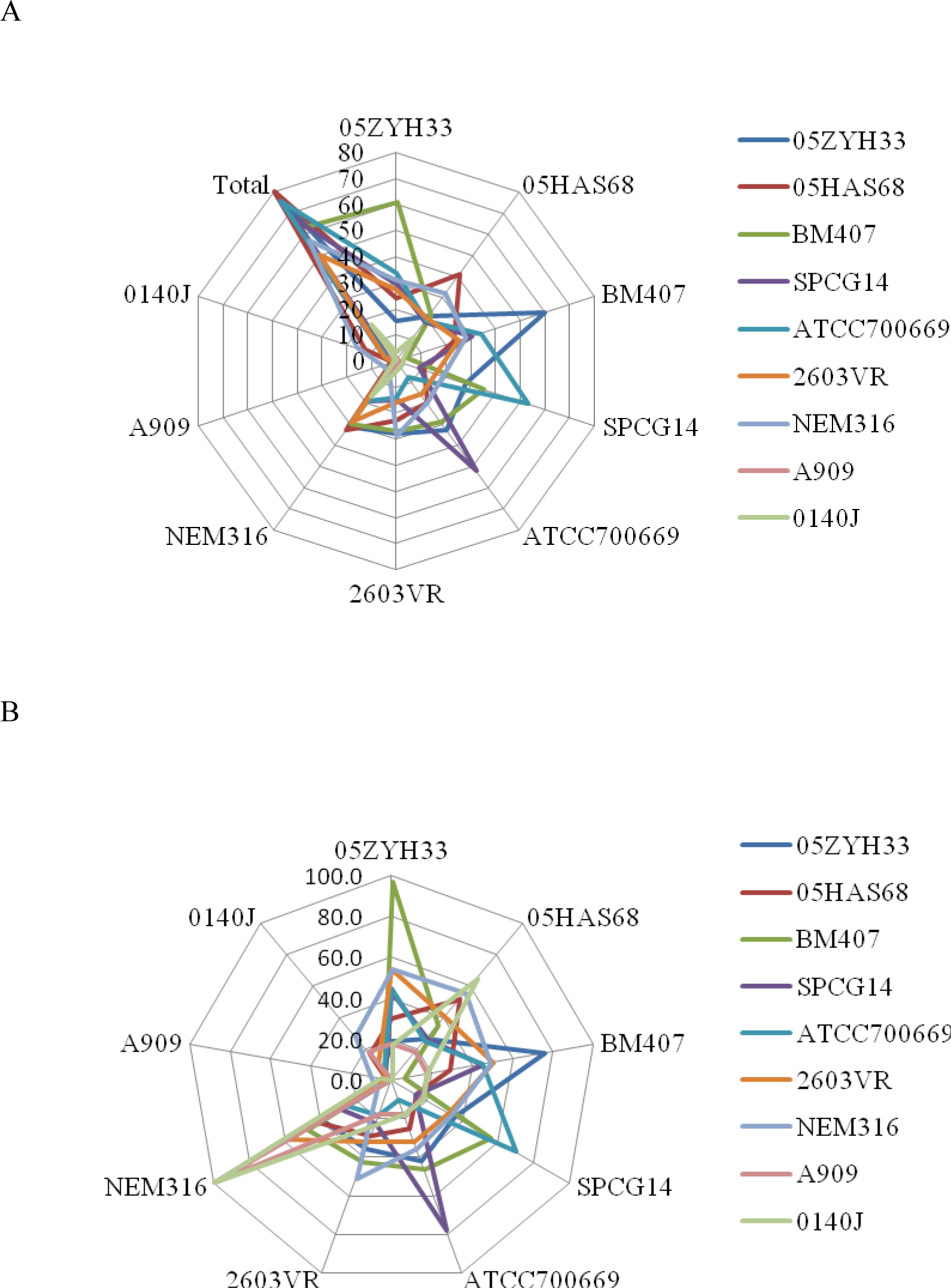
Radar diagram illustrating the distributions of homologous genes of 15SGIs. The number (A) and percentage (B) of the distributions of homologous genes were indicated in each 15SGI of 9 *Streptococcus* strains.

### Streptococcus suis

In a previous study, our group identified a novel PAI-like DNA segment of ∼89 kb in length (15SGI) as being present in the Chinese virulent strains 98HAH12 and 05ZYH33 of *Streptococcus suis.* Meanwhile, strain BM407 is a recent human clinical isolate from the patient with meningitis and was sequenced in collaboration with Dr. Constance Schultz of the Oxford University Clinical Research Unit/Hospital for Tropical Diseases and Prof. Duncan Maskell of the Centre for Veterinary Science, Ho Chi Minh City, Vietnam (Holden *et al.* 2009). The resulting 15SGIs were found to consist of 79 major open reading frames (ORFs), 05SSU0903 to 05SSU0981, and 63 major ORFs for the genomes of strains 05ZYH33 and BM407, respectively. The data revealed the presence of 12 strain-15SGI specific genes for strain 05ZYH33 and 8 genes for strain BM407. All of the genes in the 15SGI of strain BM407 were included in the 15SGI of strain 05ZYH33, except its unique genes. Notably, there were two two-component signal transduction system(TCSTS), one termed the SalK/SalR regulatory system, which is requisite for the full virulence of ethnic Chinese isolates of highly pathogenic *Streptococcus suis* serotype 2, but it was absent in the 15SGIs of the other strains (Chen *et al*. 2007). Although Holden suggested that the presence of these genes cannot be used as a marker of virulence (Holden *et al*. 2009), it is maybe keys which strain 05ZYH33 has higher virulence than other virulent strains of *Streptococcus suis* serotype 2, that has been implicated by us in an animal model (Li *et al.* 2008). The 15SGI of avirulence strain 05HAS68 consists of two tandem metabolism GIs based upon the structural features and the functional role of a majority of its genes. Two different sugar metabolism-related gene clusters emerged from the two tandem 15SGIs, respectively. They are fucose and lactose metabolic system as the novel or enhance sole carbon and energy source for growth. They share similar structural features consisting of four parts, including sugar metabolism-related enzyme gene clusters, transcriptional regulators, site-specific DNA methylase genes and a type IV secretory pathway. The fucose metabolic system is unique to strain 05HAS68. In addition, these GIs all carry one or two tetracycline-resistant genes (05SSU0922, 68SSU02364, SSUBM407_0983/0954 and SSUBM407_0954). Moreover, a toxin-antitoxin (TA) system, which was initially discovered on low copy number plasmids to ensure their segregation stability (Khoo *et al.* 2007; Li *et al.* 2011), emerged in the 15SGIs of three strains (05SSU0936, 05SSU0937; SSUBM407_0966, SSUBM407_0967 and 68SSU_02324, 68SSU_02325).

### Streptococcus agalactiae

The study identified the 15SGIs of strain 2603V/R, NEM316 and A909 of *Streptococcus agalactiae,* carrying 50, 57 and 6 genes that include 5, 16 and 0 strain-specific genes, respectively. All of the ORFs of the 15SGI of strain A909 share highly similar sets of genes (Identity >99%) in the 5’ region, which is the first 15SGI of strain NEM316. These ORFs code hypothetical protein, except site-specific recombinase (integrase, SAK_1326). The 15SGI belongs to an unknown functional GI. The 3’ region of the 15SGI of strain NEM316 appeared in a highly conserved region, which is homologous with the region of 15SGI from strain 2603VR and two virulent strains, BM407 and 05ZYH33, of *Streptococcus suis* (Fig. 2).

### Streptococcus pneumonia

*Streptococcus pneumoniae* serotype 14 isolate CGSP14 from a child with necrotizing pneumonia and *Streptococcus pneumoniae* serotype 23F isolate ATCC700669 from a nasopharyngeal colonization of a patient share features in common with multi-drug-resistant pandemic clones recently isolated. Previous study showed that CGSP14 shared the largest number of orthologous genes with strains Spn23F used for comparative genomic analysis, indicating that the CGSP14 genome shows highest homology to the sequenced strains (Ding *et al.* 2009; Croucher *et al.* 2009). The analysis of the 15SGIs gene content of strain CGSP14 and ATCC700669, consisting of 67 genes and 76 genes, respectively, show that they belong to unique number only is 3 and 4 (Fig. 3A), 13.4% and 10.5% of total genes (Fig 3B) separately. 77.6% (52/67) and 69.7% (53/76) belong to orthologous genes between the two 15SGIs. Only 19 and 31 are functionally annotated or inferred, and the remaining genes encode hypothetical proteins or uncharacterized proteins in the 15SGIs gene content of strain CGSP14 and ATCC700669, respectively. The presence of a large number of hypothetical genes conserved across the 15SGIs of these two strains suggested important roles for the putative functions of the 15SGIs. In addition, the 15SGI of strain Spn23F, which is similar to the strains of *Streptococcus suis* noted above, carries a gene encoding the tetracycline-resistance protein spn23F_13050. Both strains contained on 15SGIs are the AT system of which includes zeta toxin and epsilon antitoxin (spn23F_12620, spn23F_12630 and SPCG_1295, SPCG_1296), just like the 15SGIs of *Streptococcus suis.* The AT system has previously described that locus as part of a 27-kb pathogenicity island termed pneumococcus pathogenicity island 1 (PPI1) (Brown et al. 2001; Meinhart *et al.* 2003).

### Streptococcus uberis

*Streptococcus uberis* was reported that can utilize nutritional flexibility derived from a diversity of metabolic options to successfully occupy a discrete ecological niche (Ward *et al.* 2009). Our results revealed that the 15SGI of *Streptococcus uberis*0140J contained an eight-gene cluster lactose metabolism system highly homologous with the region of 15SGI of strain 05HAS68 and NEM316 coding genes of lactose metabolism and possibly acquired from strain 05HAS68 or NEM316 loci between the two genes coding integrase and replication initiator protein (Fig. 2). However, the lactose metabolism system of 15SGI in strain 05HAS68 is different by the absence of transcriptional regulators, site-specific DNA methylase genes and the type IV secretary pathway.

### Site-specific recombinases

Phage integrase or recombinase is the prototypical member of a large family of enzymes that catalyze site-specific DNA recombination via single-strand cleavage. They are required for the site-specific integration and excision of the GIs. We performed phylogenetic analyses of a wide collection of integrases from 9 streptococcus strains. Eleven unique integrase proteins from 9 15SGIs of *Streptococcus* were demonstrated that were more closely related to each other than others (Fig 4). In additional, we preformed BLASP and conserved domains searches from CCD database of NCBI (Marchler-Bauer *et al.* 2009) showed that the integrases were specific hits to phiLC3 phage and phage-related integrases, which belong to the tyrosine family of site-specific recombinases (Fig. S1B). They display identity scores =74%. Tyrosine recombinases mediate a wide range of important genetic rearrangement reactions or participate in diverse biological processes by catalyzing recombination between specific DNA sites. Models for tyrosine recombinases have been based largely on work done and described for the catalytic domains on the integrase of phage lambda and recombinases like Cre, Flp, and XerC/D (Malanowska et al. 2009; Bregu *et al.* 2002). Our results also showed that the 15SGI integrases share a similar region of lambda integrase-like catalytic core (Table 3). Although these integrases have high identity or similarity with each other based on their amino acids (Fig. S1A), unlike sequences are associated with other integrase families. The output from SWISS-MODEL Workspace, an automated protein homology-modeling server for the catalytic core regions, resembled the structure of site-specific recombinase IntI4 (2a3v) and site-specific recombinase Xerd (1a0p), with the conservation of important 4-helix structural elements (Fig. 5A, B). In these regions, two DNA binding regions were predicted (Fig. 5A) and a high hydrophobicity score residues located in the second helix (Fig. 5C).

**Table 3.** HMM sequence search results of site-specific integrases from 15SGIs.

| Protein | Region | Active<br>site<br>tyrosine<br>(Y) | E-value |  |
| --- | --- | --- | --- | --- |
|  |  |  | Superfamily | Family |
|  |  |  | DNA<br>breaking-rejoining<br>enzymes | Lamda<br>integrase-like,<br>catalytic core |
| 05SSU903 | 186-370 | 366 | 3.92E-33 | 0.0022 |
| 68SSU02186 | 178-360 | 358 | 1.32E-26 | 0.0035 |
| 68SSU02282 | 182-366 | 362 | 2.46E-30 | 0.002 |
| BM407_1003 | 186-370 | 366 | 8.75E-33 | 0.0022 |
| SUB0805 | 131-315 | 311 | 3.32E-30 | 0.0018 |
| SPCG1274 | 181-367 | 361 | 1.54E-35 | 0.0023 |
| SPN23F12410 | 181-367 | 361 | 1.54E-35 | 0.0026 |
| SAG1247 | 182-366 | 362 | 4.45E-33 | 0.0026 |
| gbs1314 | 179-367 | 359 | 1.29E-32 | 0.0055 |
| gbs1325 | 182-366 | 362 | 1.18E-29 | 0.0025 |
| SAK1326 | 179-367 | 359 | 1.29E-32 | 0.0055 |

**Figure 4.**
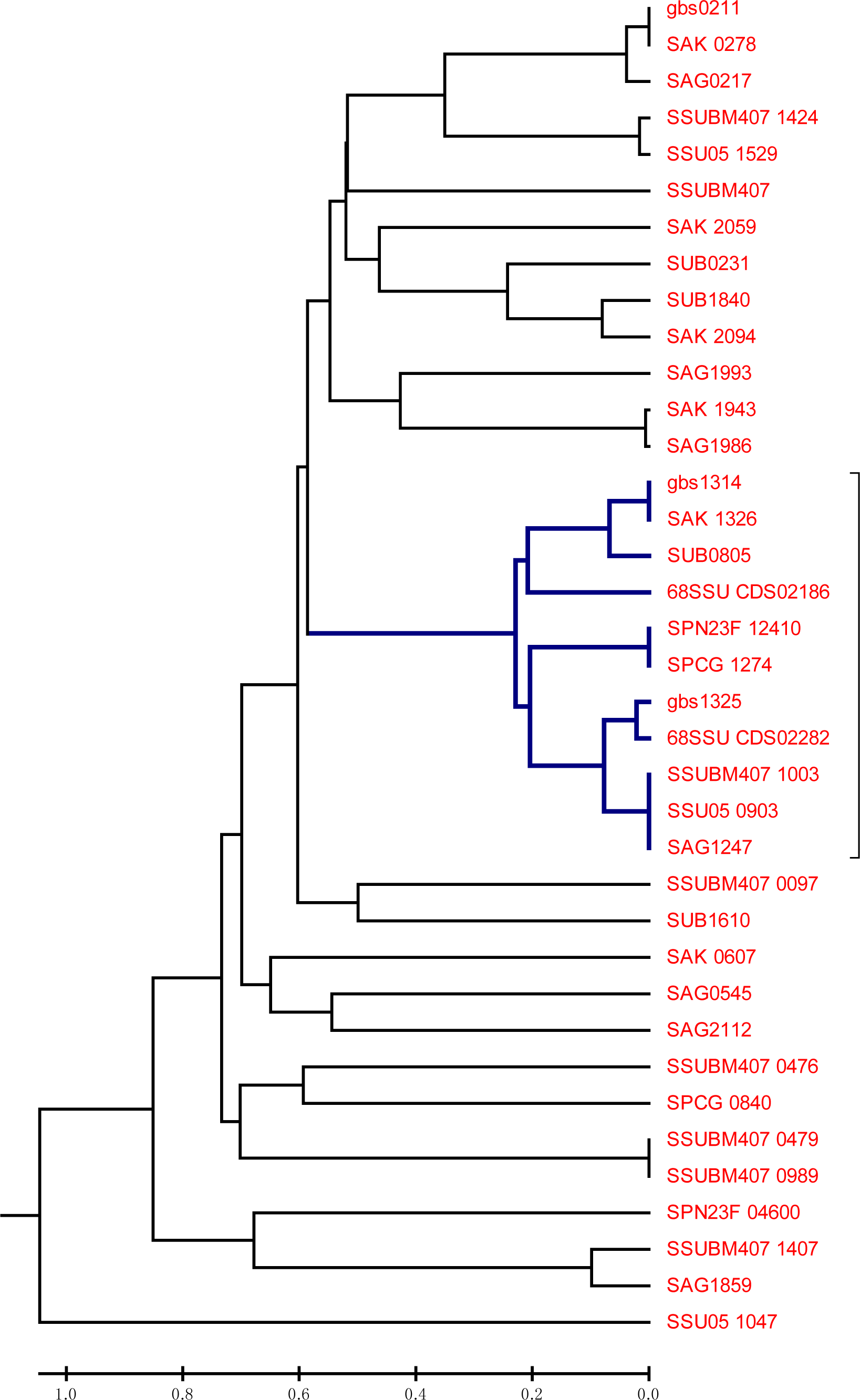
Evolutionary relationships of integrases and replicate initiator factors. The trees were constructed with integrase sequences from 9 *Streptococcus* strains. Bootstrap proportions are obtained from 500 replicates and are labeled as percentages near each internal node.

**Figure 5.**
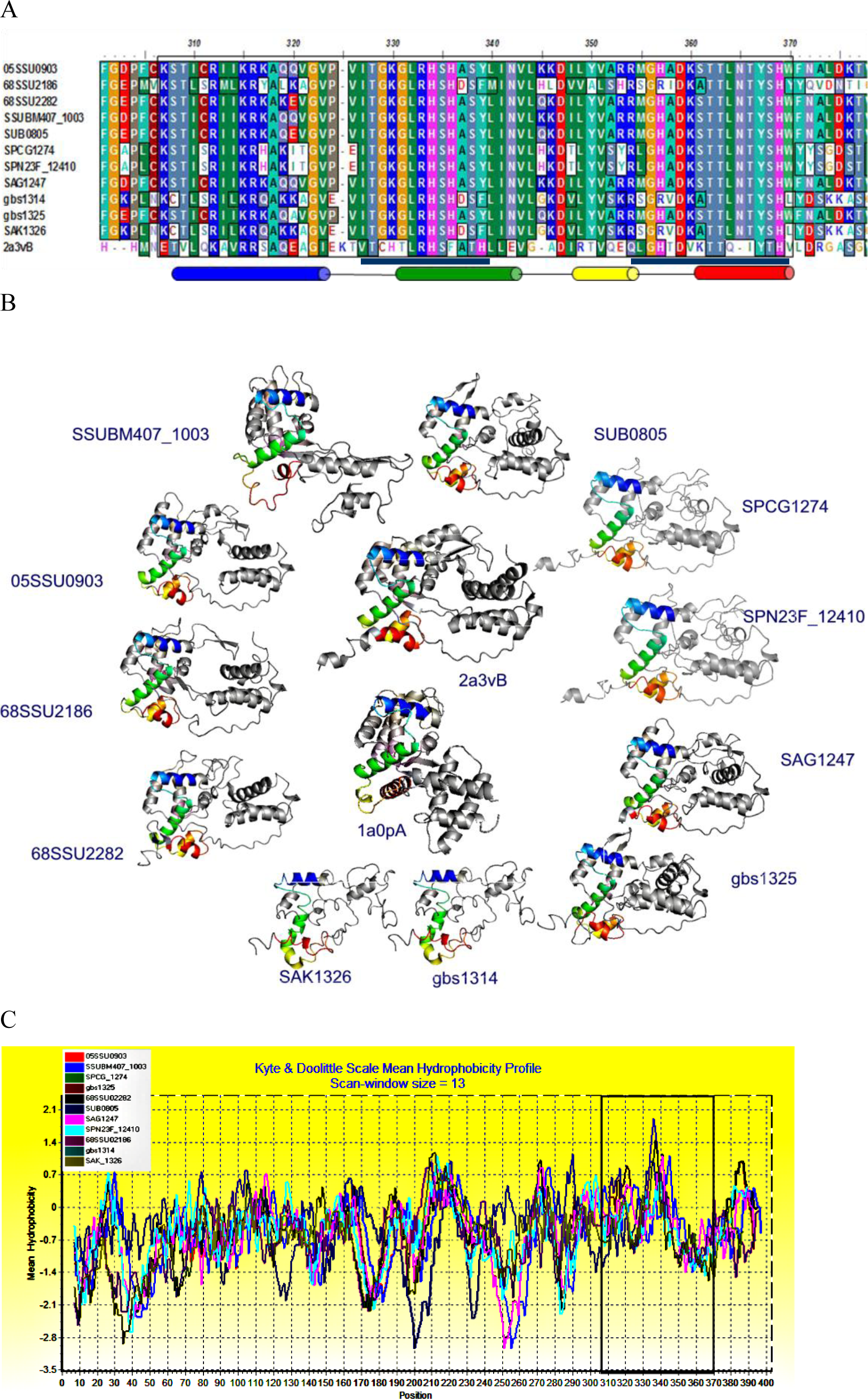
Analysis of conserved region of 11 integrases of 15SGIs. A. Multiple sequence alignment of the potential catalytic domains of a new IntI4-like integrase family from 9 15SGIs of *Streptococcus*; B. The alignment was assembled from individual pairwise alignments to IntI4 (PDB ID: 2a3v) or XerD (PDB ID: 1a0p) by PyMOL. The color structure indicated the location of conserved acid residues of each integrase. C. In this conserved region (black box), hydrophobic residues predicted by using BioEdit can be observed.

## Discussion

Our previous studies indicated that a putative pathogenicity island (PAI), termed 89k, emerged in *Streptococcus suis* serotype 2 strains 05ZYH33 and 98HAH33. The 89k GIs are flanked at their boundaries by direct repeats having a length of 15bp and a G+C content which differs significantly from that of the host organism. Therefore, starting with *a priori* assumptions about the structure of GIs, we undertook a study to resolve the structural features of GIs across the boundaries of different *Streptococcus* subspecies, using the model provided by Vernikos and Parkhill (Vernikos *et al.* 2008).

In generally speaking, the genomic positions of these GIs were not random. There are 75% of currently known genomic islands are inserted at the 3’ end of a tRNA locus. A few other genes may also act as integration sites for GIs, such as the *Salmonella*genomic island 1 (SGI1), which exhibits a site-specific integration into the *Streptococcus enterica, Escherichia coli,* or *Proteus mirabilis* chromosome at the 3' end of the *thdF* gene (Doublet *et al.* 2008). The sequence of 15bp direct repeats from the 3’ end sequences of the highly conserved ribosome *l7/l12* genes are predicted to be involved in the integration of all 15SGIs, except the 15SGI of *Streptococcus uberia*strain 0140J. To our knowledge, this l7/l12-specific recombination is reported here for the first time. The last 15 bases of the 3’ end sequences of ribosome *l7/l12* gene of *Streptococcus uberia* is TTATTTAAGAGTGAT, which differs from the TTATTTAAGAGTAAC of *Streptococcus suis, Streptococcus pneumoniae,* and *Streptococcus agalactiae*, but is identical to *Streptococcus pyogenes* and *Streptococcus equi subsp.* Based on these findings, we previously inferred that 15SGI could not integrate into the chromosome of either *Streptococcus pyogenes* or *Streptococcus equi subsp* by the difference inherent in these two key bases. However, even though the 15SGI of *Streptococcus uberia* strain 0140J is independent of the ribosome *l7/l12* gene, it adjoins a putative GI (Ward *et al.* 2009), which is inserted adjacent to the *l7/l12* gene loci. However, the general criteria previously suggested by Vernikos and Parkhill (Vernikos *et al.* 2008) do not allow us to elucidate the mechanism of acquisition.

In addition, Many structure-function relationships were discussed about site-specific recombinases (integrases) by identification of residues and secondary structures implicated in catalysis, specific and nonspecific DNA binding, protein-protein interactions and overall protein folding (Nunes-Düby et al. 1998; Douglas *et al.* 2006). Still, a full and unambiguous understanding of site-specific recombination of the 15SGI integrase family requires more solid biochemical investigation. In sum, the sequence alignments and prediction structure of the catalytic domains presented here will help guide and interpret future biochemical analyses of the 15SGIs family of integrases.

In conclusion, we employed a comparative genomics approach to define a novel GI in the genus *Streptococcus* which also appears across strains of the same species according to a previously described GI model: an 89-kb PAI-like structure of *Streptococcus suis* strain 05ZYH33 required for regulating full virulence in piglets (Chen *et al.* 2007; Li *et al.* 2008). While the features of this novel GI is in general agreement with those of the classical model for predictive GIs, marked differences in insertion stand in contrast to most tRNA genes. These GIs localize in a 15bp hotspot at the 3’ end of the l7/l10 gene by an IntI4-like integrase family. There is striking variation in gene contents, but few unique genes arise from these 15SGIs. The GIs of *Streptococcus* can be seen as a family of mobile elements with a well-defined core and variable structural features. This GI family, which was acquired by large insert of horizontally acquired DNA, seems to be of importance in species differentiation and adaption to new hosts, while it plays an important role during strain evolution in the genus *Streptococcus.*

## Supporting information

Figure S1 multiple sequence alignment (A) and conserved domains (B) on 15SGI integrase family

Figure S2 multiple sequence alignment of 15SGI replicator initiator A protein family

## Acknowledgments

We thank Fuquan Hu, Li Ming from Department of Microbiology, Third Military Medical University, Chongqing, China and Xiaoning Wang from Provincial Key laboratory of Biotechnology, School of Biosciences and Engineering, South China University of Technology, Guangzhou, China making valuable suggestions on the comparative analysis of genomic islands. This work was supported by grants to J.T. from the National Natural Science Foundation of China (#81171527 and 81371768), and to J.W. from the Key Issue of Medical and Health Foundation of Nanjing Military Command (#12Z17 and 09Z015)..

## Authors’ contributions

J.T., C.J., Y.F., S.J. and J.W.conceived and designed the experiment; J.W., Y.Y. and W.F. performed the comparative genomics and analyzed date. L.Z., L.L. and J.W. participated in the genomic island identification. Y.F., S.J. JW. and JT. prepared the manuscript. All authors read and approved the final manuscript.

## References

Arnold K, Bordoli L, Kopp J, Schwede T. 2006 The SWISS-MODEL Workspace: A web-based environment for protein structure homology modelling. Bioinformatics 22: 195–201.

Bregu M, Sherratt DJ, Colloms SD. 2002 Accessory factors determine the order of strand exchange in Xer recombination at psi. EMBO J. 21: 3888–3897.

Brown JS, Gilliland SM, Holden DW. 2001 A streptococcus pneumonilea pathogenicity island encoding an ABC transporter involved in iron uptake and virulence. Mol. Microbiol. 40, 572–585

Brown JS, Gilliland SM, Spratt BG, Holden DW. 2004 A locus contained within a variable region of pneumococcus pathogenicity island 1 contributed to virulence in mice. Infect Immun 72: 1587–1593.

Chen C, Tang J, Dong W, Wang C, Feng Y, et al. 2007 A glimpse of streptococcus toxic shock syndrome from comparative genomics of S. suis 2 Chinese isolates. PLoS One 2: e315.

Croucher NJ, Walker D, Romero P, Lennard N, Paterson GK, et al. 2009 Role of conjugative elements in the evolution of the multidrug-resistant pandemic clone Streptococcus pneumonia Spain23F ST81. J. Bacteriol 191: 1480–1489.

Ding F, Tang P, Hsu MH, Cui P, Hu S, et al. 2009 Genome evolution driven by host adaptations results in a more virulent and antimicrobial-resistant Streptococcus pneumoniae serotype 14. BMC Genomics 10:158.

Dmitriev A, Yang YH, Shen AD, Totolian A. 2006 Adjacent location of the bac gene and two-component regulatory system genes within the putative Streptococcus agalactiae pathogenicity island. Folia Microbiol Praha 51: 229–235

Dobrindt U, Hochhut B, Hentschel U, Hacker J. 2004 Genomic islands in pathogenic and environmental microorganisms. Nat Rev Microbiol. 2: 414–424.

Doublet B, Golding GR, Mulvey MR, Cloeckaert A. 2008 Secondary chromosomal attachment site and tandem integration of mobilizable salmonella genomic island 1. PLoS One 3: e2060.

Douglas MD, Gaëlle D, Marie B, Didier M, Deshmukh NG 2006 Structural basis for broad DNA-specificity in integron recombination. Nature 440: 1157–1162.

Glaser P, Rusniok C, Buchrieser C, Chevalier F, Frangeul L, et al. 2002 Genome sequence of Streptococcus agalactiae, a pathogen causing invasive neonatal disease. Mol. Microbiol 45: 1499–1513

Herbert MA, Beveridge CJ, McCormick D, Aten E, Jones N, et al. 2005 Genetic islands of Streptococcus agalactiae strains NEM316 and 2603VR and their presence in other Group B streptococcus strains. BMC Microbiol 5:31.

Holden MT, Hauser H, Sanders M, Ngo TH, Cherevach I, et al. 2009 Rapid evolution of virulence and drug resistance in the emerging zoonotic pathogen Streptococcus suis. PLoS One 4: e6072

Juhas M, van der Meer JR, Gaillard M, Harding RM, Hood DW, et al. 2009 Genomic islands: tools of bacterial horizontal gene transfer and evolution. FEMS Microbiol Rev 33: 376–393.

Khoo SK, Loll B, Chan WT, Shoeman RL, Ngoo L, et al. 2007 Molecular and structural characterization of the PezAT chromosomal toxin-antitoxin system of the human pathogen Streptococcus pneumoniae. J Biol Chem 282: 19606–19618.

Lefébure T, Stanhope MJ 2007 Evolution of the core and pan-genome of Streptococcus: positive selection, recombination, and genome composition. Genome Biol 8:R71.

Li M, Shen X, Yan J, Han H, Zheng B, et al. 2011 GI-type T4SS-mediated horizontal transfer of the 89K pathogenicity island in epidemic Streptococcus suis serotype 2. Mol Microbiol. 79: 1670–1683.

Li M, Wang C, Feng Y, Pan X, Cheng G et al. 2008 SalK/SalR, a two-component signal transduction system, is essential for full virulence of highly invasive Streptococcus suis serotype 2. PLoS One 3: e2080.

Malanowska K, Cioni J, Swalla BM, Salyers A, Gardner JF. 2009 Mutational analysis and homology-based modeling of the IntDOT core-binding domain. J Bacteriol 191: 2330–2339.

Marchler-Bauer A, Anderson JB, Chitsaz F, Derbyshire MK, DeWeese-Scott C, et al. 2009 CDD: specific functional annotation with the Conserved Domain Database. Nucleic Acids Res 37: D205–210.

Marri PR, Hao W, Golding G.B 2006 Gene gain and gene loss in streptococcus: is it driven by habitat? Mol. Biol. Evol. 23: 2379–2391

Meinhart A, Alonso JC, Sträter N, Saenger W. 2003 Crystal structure of the plasmid maintenance system epsilon/zeta: functional mechanism of toxin zeta and inactivation by epsilon 2 zeta 2 complex formation. Proc Natl Acad Sci U S A 100: 1661–1666.

Nunes-Düby SE, Kwon HJ, Tirumalai RS, Ellenberger T, Landy A. 1998 Similarities and differences among 105 members of the Int family of site-specific recombinases. Nucleic Acids Res. 26: 391–406.

Petzer IM, Karzis J, Watermeyer JC, van der Schans TJ, van Reenen R. 2009 Trends in udder health and emerging mastitogenic pathogens in South African dairy herds. J S Afr Vet Assoc 80: 17–22.

Rosini R, Rinaudo CD, Soriani M, Lauer P, Mora M, et al. 2006 Identification of novel genomic islands coding for antigenic pilus-like structures in Streptococcus agalactiae. Mol Microbiol. 61: 126–141.

Sentchilo V, Czechowska K, Pradervand N, Minoia M, Miyazaki R, et al. 2009 Intracellular excision and reintegration dynamics of the ICEclc genomic island of Pseudomonas knackmussii sp. strain B13. Mol Microbiol 72: 1293–1306.

Tamura K, Dudley J, Nei M, Kumar S. 2007 MEGA4: Molecular Evolutionary Genetics Analysis MEGA software version 4.0. Molecular Biology and Evolution 24: 1596–1599.

Tuanyok A, Leadem BR, Auerbach RK, Beckstrom-Sternberg SM, Beckstrom-Sternberg JS, et al. 2008 Genomic islands from five strains of Burkholderia pseudomallei. BMC Genomics. 9: 5666.

Vernikos GS, Parkhill J. 2008 Resolving the structural features of genomic islands: A machine learning approach. Genome Res 18: 331–342.

Ward PN, Holden MTG, Leigh JA, Lennard N, Bignell A, et al. 2009 Evidence for niche adaptation in the genome of the bovine pathogen Streptococcus uberis. BMC Genet 10: 54.

Zhao, Y., Liu G, Li S, Wang M, Song J, et al. 2011 Role of a type IV-like secretion system of Streptococcus suis 2 in the development of streptococcal toxic shock syndrome. J Infect Dis. 204, 274–281.

